# Engineering a more specific *E. coli* glyoxylate/hydroxypyruvate reductase for coupled steady state kinetics assays

**DOI:** 10.1101/2022.04.02.486822

**Authors:** Nemanja Vuksanovic, Dante A. Serrano, Brandon M. Patterson, Nicholas R. Silvaggi

## Abstract

The *E. coli* glyoxylate reductase/hydroxypyruvate reductase A (EcGhrA) was investigated as a coupling enzyme to monitor the transamination of 2-ketoarginine and glycine by the L-enduracididine biosynthetic enzyme MppQ. Surprisingly, 2-ketoarginine proved to be an efficient substrate for EcGhrA. Since the promiscuity of EcGhrA prevented its use as a coupling enzyme to monitor the aminotransferase activity of MppQ, we set about engineering a more specific variant. X-ray crystal structures of EcGhrA were determined in the unliganded state, as well as with glyoxylate and 2-ketoarginine bound. The electron density maps of EcGhrA with 2-ketoarginine bound showed weak electron density for the side chain of this substrate, complicating the choice of active site residues to target for site-directed mutagenesis. The structure of the complex did, however, suggest that the side chain of W45 could interact with the guanidinium group of 2-ketoarginine. We therefore generated the EcGhrA^W45F^ variant and tested it for activity with 2-ketoarginine, glyoxylate, oxaloacetate, α-ketoglutarate, α-oxofuranacetic acid, phenyl pyruvate, 3-mercaptopyruvate and 2-ketobutyric acid. The W45F variant exhibited a ∼10-fold decrease in the specificity constant (k_cat_/K_M_) for 2-ketoarginine, while the reaction with glyoxylate was not significantly impaired. The reactions of the W45F variant with the alternative substrates oxaloacetate and α-ketoglutarate were also impaired. Thus, the W45F variant is a less promiscuous enzyme than the wild-type. This engineered EcGhrA^W45F^ variant could be generally useful as a coupling system for enzymes that produce glyoxylate, such as 4-hydroxy-2-oxoglutarate aldolase or isocitrate lyase.

---

The non-proteinogenic amino acid L-enduracididine (Scheme 1, **5**) is a component of several non-ribosomally produced peptide natural products, including the antibiotic mannopeptimycin ^*1,2*^. As a part of our work on L-enduracididine biosynthesis,^*3-5*^ it became necessary to develop a coupled assay to measure the kinetics of the putative 2-ketoenduracididine aminotransferase MppQ. In addition to its proposed role in the last step of L-enduracididine biosynthesis, MppQ is also capable of returning the dead-end product 2-ketoarginine (Scheme 1, **3**) back to the starting material, L-Arg. Preliminary experiments showed that, in reactions with L-Arg and a selection of amino acceptor substrates (*e.g*. glyoxylate, pyruvate, oxaloacetate and α-ketoglutarate, Scheme 2), MppQ was most efficient using glyoxylate as the acceptor substrate (unpublished data). This led to the idea that, in the more biosynthetically relevant transamination of 2-ketoenduracididine or 2-ketoarginine to the corresponding amino acids, glycine should be an efficient amino donor substrate. Since these reactions would result in the production of glyoxylate, we selected GhrA from *E. coli* (EcGhrA) as a potential coupling enzyme in a coupled spectrophotometric assay for the transamination of 2-ketoarginine or 2-ketoenduracididine and glycine. This particular enzyme was chosen over other enzymes previously used in assays of glyoxylate-producing reactions, such as porcine L-lactate dehydrogenase^*6*^ or the glyoxylate reductase from *Spinacea olerac*ea,^*7*^ because it could be produced very efficiently and inexpensively in-house.

**Scheme 1.**
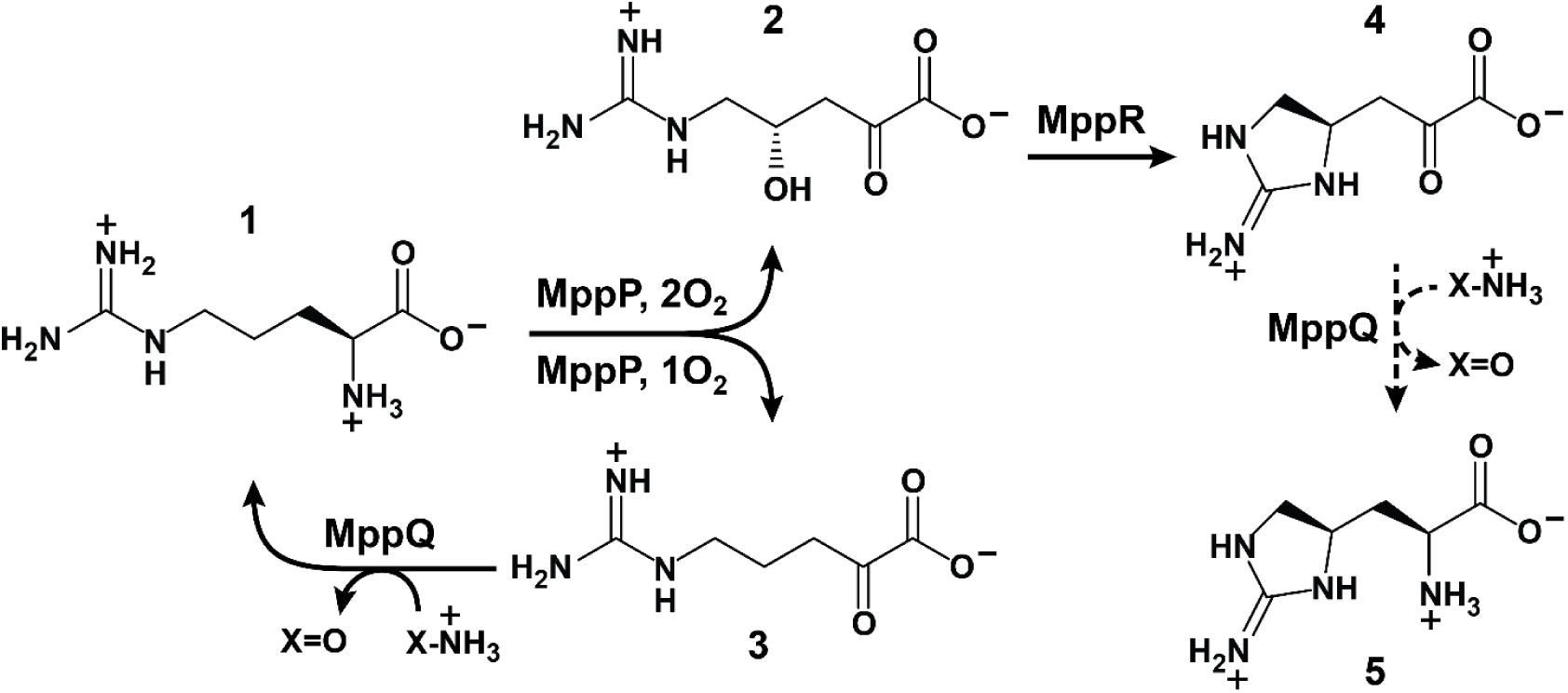
The L-enduracididine biosynthetic pathway begins with L-arginine (**1**), which is oxidized by the unusual PLP-dependent L-Arg oxidase, MppP, to 4(S)-hydroxy-2-ketoarginine (**2**). MppP, at least *in vitro*, releases the partially oxidized dead-end product 2-ketoarginine (**3**) in about 40% of catalytic cycles. The oxidized arginine derivative **2** is cyclized by MppR to create the iminoimidazolidine ring of the intermediate **4** (*i.e*. 2-ketoenduracididine). MppQ is thought to catalyze the transamination of **4** with an as-yet-unidentified amino donor substrate to yield the finished amino acid L-enduracididine (**5**). MppQ also catalyzes the transamination of the dead-end product **3** to regenerate L-Arg, though it has not been ascertained if this is a physiologically relevant catalytic activity or an adventitious *in vitro* phenomenon.

The D-2-hydroxyacid dehydrogenases (2HAD) family contains enzymes with diverse substrate specificities that show significant potential for use in the industrial synthesis of chiral D-2-hydroxy acids.^*8*^ Members of the 2HAD family are known to be promiscuous, and EcGhrA also catalyzes the reduction of 2-hydroxypyruvate.^*9*^ Surprisingly (at least to us), the promiscuity of EcGhrA extended to the much larger and positively charged 2-ketoarginine, which would, of course, preclude its use in monitoring the reaction we wished to study.

This observation led us to wonder what other α-keto acids are substrates for EcGhrA. The substrate specificities of other GhrA homologs have been examined and it was found that oxaloacetate is a substrate for the *Euglena gracilis*^*10*^ and *Saccharomyces cerevisiae*^*11*^ enzymes, succinic semialdehyde is a substrate for the *Arabidopsis thaliana* homolog^*12*^, *Neurospora crassa* GhrA reacts with phenylpyruvate,^*13*^ and α-ketoglutarate is a slow substrate for the *E. gracilis* GhrA^*10*^. To extend the knowledge of EcGhrA promiscuity, we investigated additional α-keto acids (Scheme 2, **7-12**), some of which have not been tested previously. The results of these studies, together with the structural work described herein, suggested that it might be possible engineer an EcGhrA variant with increased specificity for glyoxylate in order to develop a suitable coupling enzyme for use in future steady state kinetics work.

**Scheme 2.**
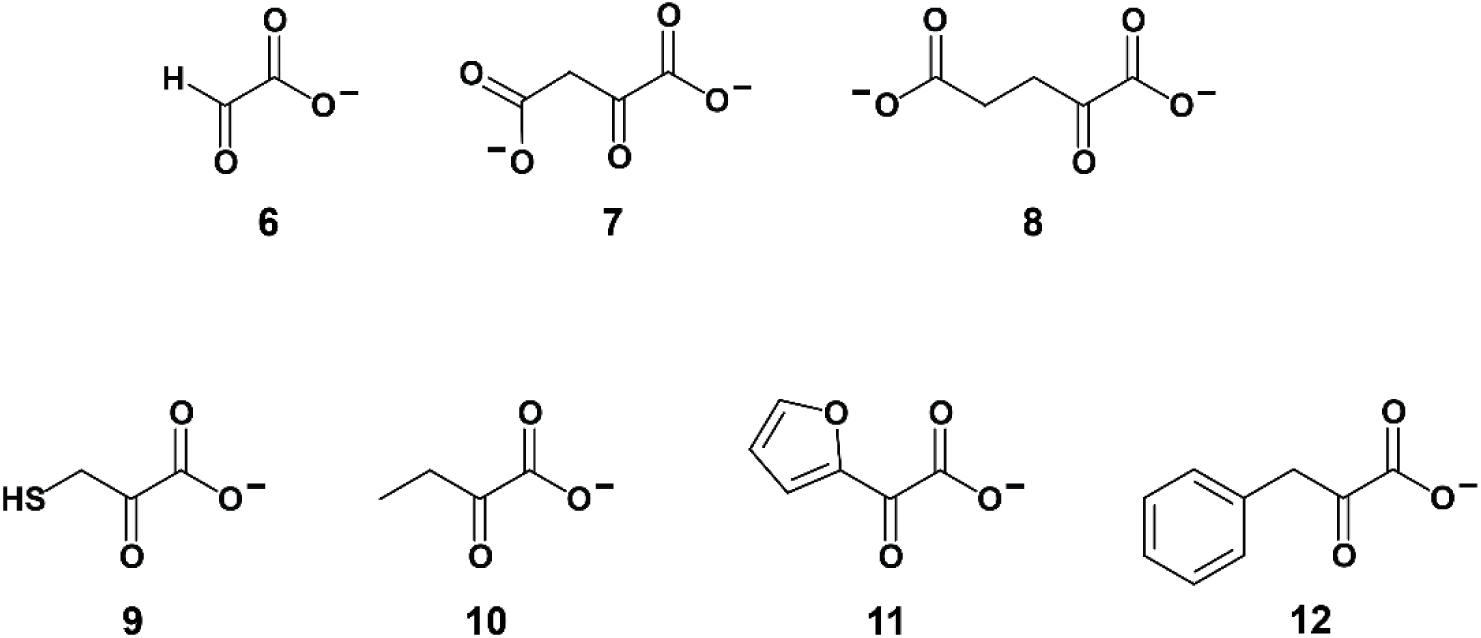
EcGhrA substrates investigated in this work.

Rational engineering of 2HAD enzymes to alter substrate specificity has been described in the literature. In 2007, Shinoda et al. transformed formate dehydrogenase into a glyoxylate reductase by mutating two catalytic residues^*14*^. In 2013, Zhen et al. rationally engineered D-lactate dehydrogenase to reduce phenylpyruvic acid, α-ketobutyric acid, α-ketovaleric acid, and α-hydroxypyruvic acid via a single Y52L mutation.^*15*^ At the time of this writing, rational engineering of a GhrA homolog to make it more specific for glyoxylate has not been attempted.

Herein we describe the crystal structures of wild-type EcGhrA in ternary complexes with NADP^+^ and either glyoxylate or **3** that were the basis of our (ir)rational design of a more specific glyoxylate reductase. These structures suggested that making a limited number of mutations near the outer edge of the active site could reduce the affinity of EcGhrA for **3**. Steady state kinetics of the wild-type and variant EcGhrA enzymes shows that a single mutation, W45F, leads to a 10-fold decrease in the specificity constant (k_cat_/K_M_) for **3** without significant impact on the reaction with glyoxylate. This new variant of EcGhrA may be useful for studying the kinetics of glyoxylate-producing enzymes where the promiscuity of the wild-type EcGhrA is problematic.

## Materials and Methods

### Cloning, expression and purification of wild-type and mutant EcGhrA

The EcGhrA gene was amplified from *E. coli* genomic DNA using primers containing NdeI and BamHI restriction sites (underlined; forward, 5’-GGATTC<u>CATATG</u>*GAAAACCTGTATTTTCAGGGT*ATGGA-TATCATCTTTTATCACCCAA-3’; reverse, 5’-CG<u>GGATCC</u>TTAGTAGCCGCGTGCGC-3’). The forward primer also encoded a tobacco etch virus (TEV) protease cleavage site (italic, translates to MENLYFQG). The PCR reaction was carried out using Phusion High Fidelity PCR Master Mix (Thermo Scientific) and the genomic DNA was extracted from *E. coli* BL21 Star (DE3) (Invitrogen) using genomic the Blood and Cell Culture DNA mini kit (Qiagen). The amplified PCR product was digested in one step using NdeI and BamHI HF (New England Biolabs) and ligated into similarly treated pET-15b to create plasmid pET15b-EcGhrA. Site-directed mutagenesis to create the EcGhrA^W45F^ variant was performed using the Q5 Site-Directed Mutagenesis Kit (New England Biolabs) using the mutagenic primers 5’-TGCTTTAGTC<u>TTT</u>CATCCTCCTG-3’ (forward) and 5’-TAATCAGCAGAGTCATTATC-3’ (reverse).

The His_6_-tagged wild-type EcGhrA was expressed from *E. coli* BL21 Star (DE3) cells carrying the pET15b-EcGhrA plasmid grown at 37 °C in Luria-Bertani broth with 100 µg/mL ampicillin. Protein expression was induced with 0.4 mM IPTG when the OD_600nm_ reached 0.6. The temperature was reduced to 25 °C and the cells were grown overnight with shaking at 250 rpm. Cells were harvested via centrifugation and resuspended in 5 mL of Buffer A [25 mM TRIS (pH 8.0), 300 mM NaCl, 10 mM imidazole] containing 0.1 mg/mL Dnase I (Worthington Biochemical Corp.) per gram of wet cell paste. Cells were lysed using a Branson Sonifier S-450 cell disruptor (Branson Ultrasonics Corp.) at 60% amplitude with 30 s pulses separated by 50 s rest periods for a total of 10 min of sonication. The temperature of the lysate was maintained below 4 °C by keeping the steel beaker suspended in an ice bath over a stir bar. The lysate was clarified by centrifugation at 39,000 x g for 45 min before loading it onto a 5 mL HisTrap column (GE Lifesciences) at a flow rate of 5 mL/min. The His_6_-tagged protein was purified using a 4-step gradient of increasing concentrations of Buffer B [25 mM TRIS (pH 8.0), 300 mM NaCl, and 250 mM imidazole; 5 column volumes each at 5, 15, 50, and 100% B]. The elution was monitored spectroscopically at 280 nm. His_6_-EcGhrA eluted at 50% Buffer B, and these fractions were collected and analysed using Coomassie-stained SDS-PAGE. In order to remove the His_6_ tag, the pooled fractions were dialyzed overnight against 10 µM TEVpM2 (an optimized TEV protease variant)^*16*^ in 50 mM TRIS (pH 8.0), 100 mM NaCl, 1 mM DTT, and 0.25 mM EDTA at 4 °C. The dialysate was passed through the same HisTrap column, but less than 10% of the protein had the His_6_ tag removed. The uncleaved protein was desalted into 10 mM HEPES pH 7.5, 1 mM DTT, by dialyzing overnight at 4 °C before being aliquoted and stored at -80 °C. EcGhrA^W45F^ was expressed and purified in the same manner as the wild type EcGhrA.

### Synthesis of 2-ketoarginine, **3**

Compound **3** was synthesized by enzymatic oxidation of L-arginine as described by Meister^*17*^ and purified as follows. Residual L-arginine was removed by passing the mixture through Dowex 50 WX8 (Sigma Aldrich) that had been equilibrated with approximately 10 CV of 0.5 M HCl, followed by 10 CV of 0.5 M NaCl, and finally 10 CV of water^*18*^. The identity and purity of **3** were confirmed by NMR spectroscopy (see Supplementary Information, Figure S1).

### Steady state enzyme kinetics

The initial velocities of the wild-type and mutant EcGhrA-catalyzed reductions of varied concentrations of glyoxylate, and concomitant oxidation of NADPH, under steady state conditions were monitored at 340 nm (ε_340nm_ = 6220 M^-1^cm^-1^) using a Shimadzu UV-2600 double beam UV-visible spectrophotometer. All the reactions were carried out in triplicate under identical conditions. The enzyme (0.5 µM) was incubated with NADPH (200 µM) for 30 seconds in 50 mM HEPES, pH 7.5 at 22 °C before initiating the reaction by the addition of substrate. The substrate concentrations ranged from 0.2 to 40 mM for **6** and **3**, and 0.6 to 40 mM for **7** and **8**. Compounds **9-12** were tested only at 40 mM. No reactions were observed with these substrates, so they were not tested at lower concentrations. The steady state kinetic constants for the reactions with evidence of substrate inhibition (*e.g*. wild-type EcGhrA with **6**) were estimated by fitting the initial velocity data to Equation 1:

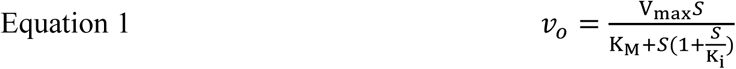

where v_o_ is the initial velocity, V_max_ is the maximum velocity, S is the substrate concentration, K_M_ is the Michaelis constant, and K_i_ is the inhibition constant for reversible substrate inhibition. Reactions showing no evidence of substrate inhibition were analyzed using the standard Michaelis-Menten equation (Equation 2):

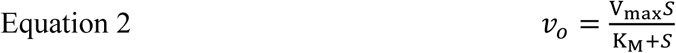

### Crystallization of the EcGhrA·NADP^+^ binary complex

Diffraction quality crystals were obtained after screening ∼16 mg/mL EcGhrA against the Index HT screen (Hampton Research). The optimized crystallization buffer consisted of 1.26 M sodium phosphate monobasic monohydrate, 0.14 M potassium phosphate dibasic, pH 5.6, and 1.8 mM NADP^+^. Hanging drop vapor diffusion experiments were initiated by mixing 1 µL of ∼16 mg/mL wild-type EcGhrA in 10 mM HEPES pH 7.5, 1 mM DTT with 1 µL of the crystallization solution on a siliconized glass cover slip and sealing this above a well containing 500 µL of the crystallization solution in a 24-well VDX plate (Hampton Research). Hexagonal prisms of approximate dimensions 200 × 50 × 50 µm formed after 1 day. For the binary complex, crystals were cryo-protected by a brief soak in the crystallization buffer supplemented with 20 % glycerol.

### Crystallization of the ternary complexes of EcGhrA with NADP^+^ and **3, 6**, or **8**

Since we observed that phosphate significantly inhibits EcGhrA activity at the concentrations used for crystallization (data not shown) and is therefore likely to compete with the substrate for binding at the active site, we identified an alternative, phosphate-free crystallization condition. After optimization of the initial condition from the Index HT screen, diffraction-quality crystals were obtained from hanging drop vapor diffusion experiments where the well solution consisted of 17% PEG 3350, 0.2 lithium sulfate, 0.1 M BIS-TRIS, pH 5.5, and 1.8 mM NADP^+^. The crystals were soaked overnight in 10 µL of a solution containing 50 mM substrate (**3, 6**, or **8**), 5 mM NADP^+^, 30% PEG 3350, 0.2 M lithium sulfate, and 0.1 M BIS-TRIS, pH 5.5. Paratone N (Hampton Research) was used to cryo-protect the crystals before being flash-cooled by plunging in liquid nitrogen.

### Crystallization of EcGhrAW45Fwith NADP+

Diffraction-quality crystals were obtained from the LMB screen (Molecular Dimensions) at 295 K, in a sitting drop vapor diffusion experiment where the well solution consisted of 16% PEG 4000, 0.1 ammonium sulfate, 0.1 M sodium citrate, pH 5.8, 20% glycerol and 1 mM NADP^+^. The crystal used for data collection was taken directly out of the drop and flash-cooled by plunging in liquid nitrogen. No further cryo-protection was required owing to the 20% glycerol present in the crystallization mixture.

### Data collection and structure solution

X-Ray diffraction data for the EcGhrA·NADP^+^ binary complex and the EcGhrA·NADP^+^·**3** ternary complex were collected using the Life Sciences Collaborative Access Team (LS-CAT) beamline 21-ID-F at the Advanced Photon Source (APS). Diffraction data for the EcGhrA·NADP^+^·**6** and EcGhrA·NADP^+^·**8** ternary complexes were collected using LS-CAT beamline 21-ID-G. Both beamlines were equipped with MAR 300 CCD detectors and 50 × 50 µm beams at wavelengths of 0.97856 Å and 0.97872 Å, respectively. In all cases, data were collected for 180 ° sweeps of the crystals with an oscillation range of 0.5° for each image. The exposure time was 1.0 s per image (2.0 s per °). The crystal-to-detector distances for the EcGhrA·NADP^*+*^ binary complex and the three ternary complexes ranged from 250 to 300 mm. The data for the EcGhrA^W45F^·NADP^*+*^ binary complex was collected on LS-CAT beamline 21-ID-D. This beamline was equipped with a Dectris Eiger 9M detector and a 50 × 50 µm beam at a wavelength of 1.1272 Å. A total of 900 frames were collected from φ = 0 to 180 ° at a scan rate of 5 °/sec, with 5 images collected per degree, at a detector distance of 200 mm. All X-ray diffraction data sets were indexed, integrated and scaled using HKL2000^*19*^.

Initial phase estimates for all five structures were determined by molecular replacement in PHASER^*20*^ as implemented in the CCP4^*21*^ suite. The structure of the binary complex of EcGhrA with NADP^+^ was determined using the coordinates of chain A of GhrA from *Salmonella typhimurium* (PDB ID 3KBO, unpublished) as the search model, with all non-protein atoms removed and all isotropic B-factors set to 20 Å^2^. After refinement of this model (see below), it was used as the search model for phasing the three ternary complex structures and the EcGhrA^W45F^ structure. The EcGhrA·NADP^+^ structure was prepared for molecular replacement by removing all hydrogen atoms, solvent molecules, and NADP^+^ and setting all isotropic B-factors to 20.0 Å^2^. Model refinement was performed using phenix.refine from the PHENIX suite^*22, 23*^ and model building was done with COOT^*24, 25*^. Iterative rounds of model building and refinement were done until the R-factors converged, at which point the substrates were modeled into the ternary complex structures. The optimal number of translation-libration-screw (TLS) groups was determined using phenix.find_tls_groups (P. V. Afonine, unpublished). The final models were validated using MolProbity^*26*^, as implemented in the PHENIX suite. Residues lacking defined electron density for the sidechain were modeled without the sidechains in order to avoid interpretational bias. The geometric restraints for **3** were calculated using phenix.elbow^*27*^. The restraints for NADP+, **6**, and **8** are distributed with the CCP4 package and were used in their respective refinements. 3-Dimensional renderings of the structures were generated using the POVSCRIPT+ modification of MOLSCRIPT^*28, 29*^. The data collection and model refinement statistics are listed in Table 1. Coordinates have been deposited in the Protein Data Bank; the accession codes for each structure are also given in Table 1.

**Table 1:**
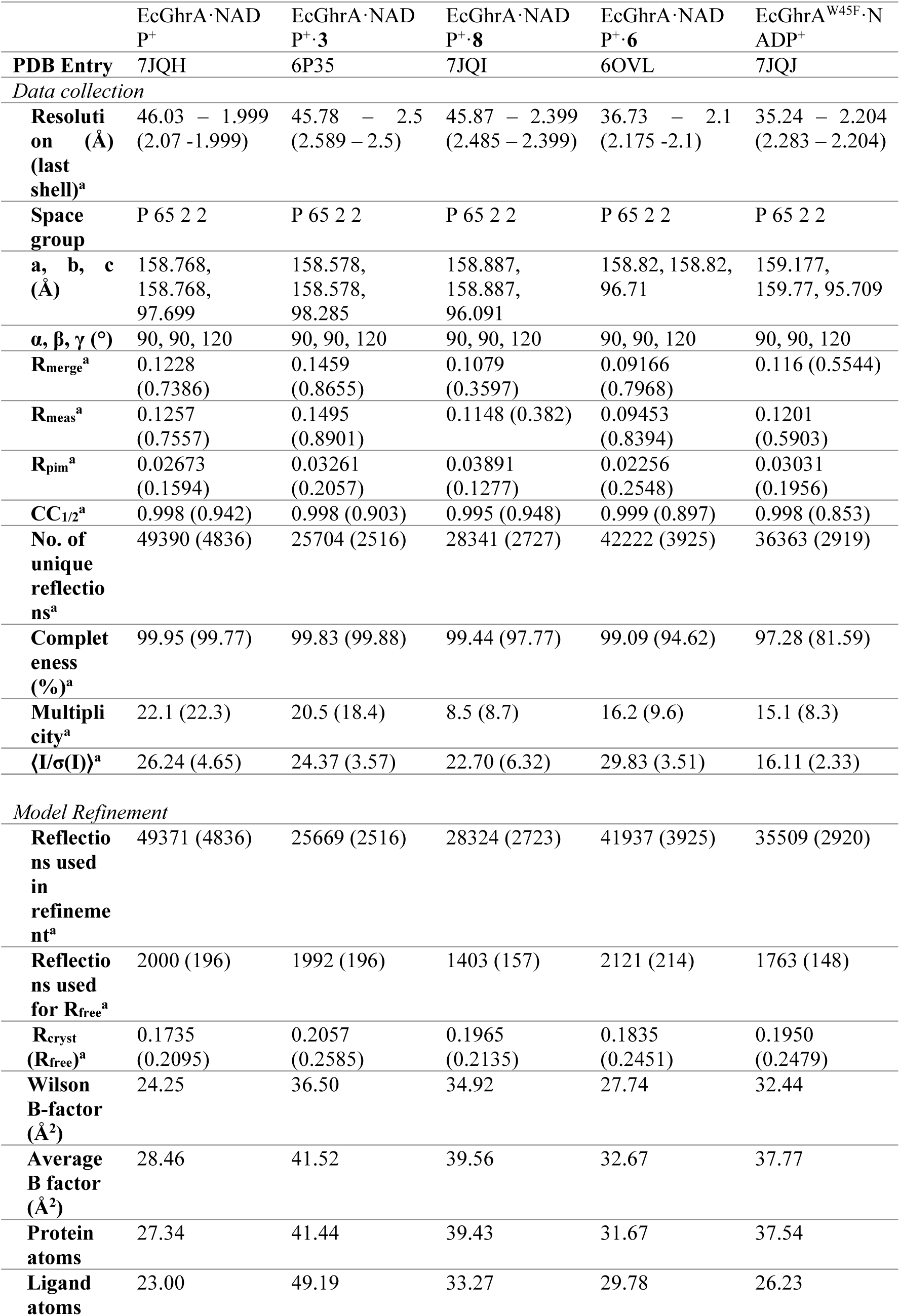
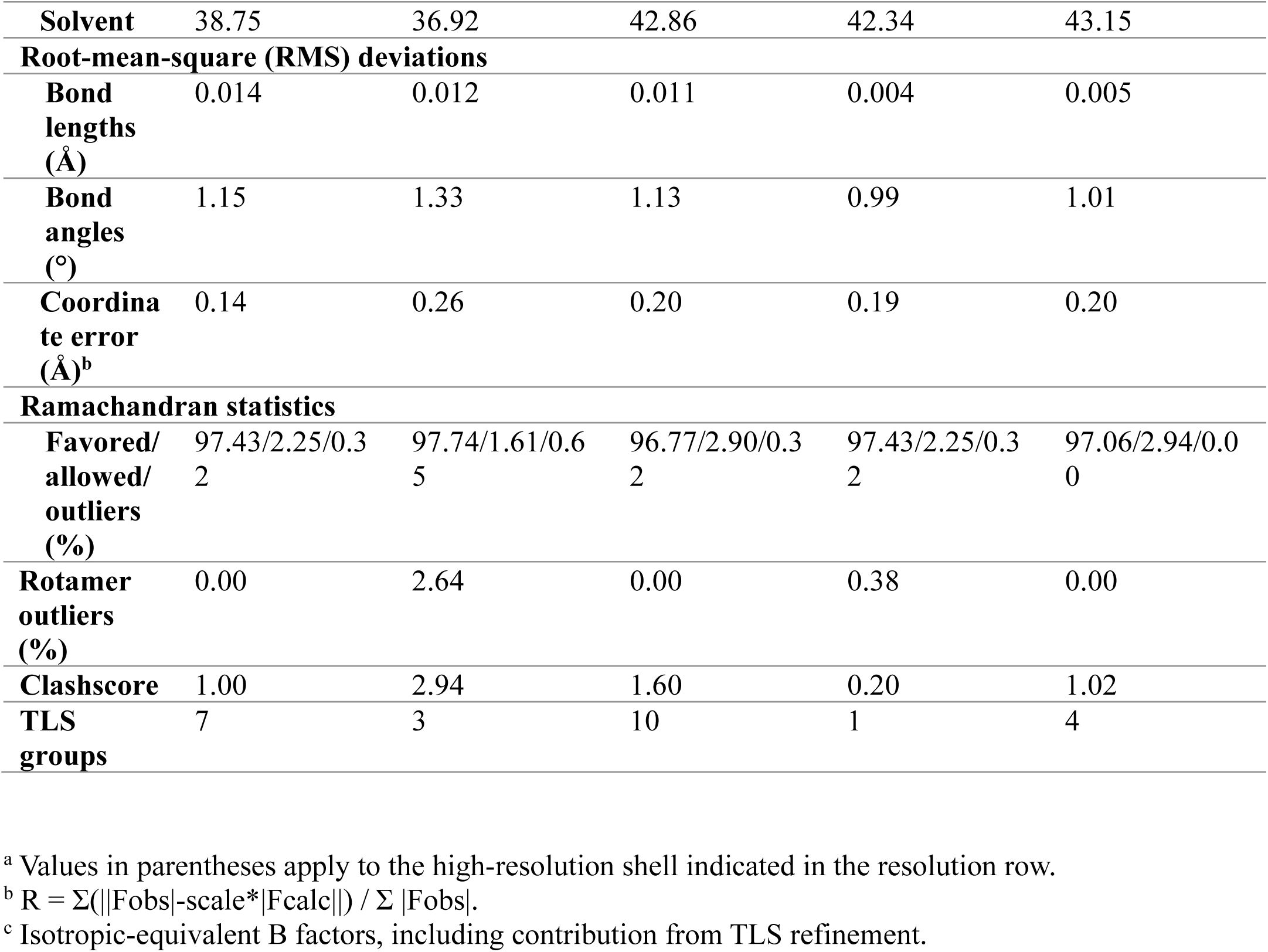
Crystallographic data collection and refinement statistics.

## Results and Discussion

### Structure determination of EcGhrA with NADP+ alone and in the presence of α-keto acid substrates

Since there were no crystal structures of the *E. coli* glyoxylate/hydroxypyruvate reductase available, we began by determining the structure of the EcGhrA·NADP^+^ complex. This form of the enzyme crystallized with two molecules per asymmetric unit arranged as a homodimer, as observed in the structures of homologous glyoxylate/hydroxypyruvate reductases^*30, 31*^. GhrA homologs from other organisms are known to be active as homodimers^*32, 33*^. Both the overall fold (Figure 1A) and active site architecture (Figure 1B) of EcGhrA are very similar to those of enzymes like *Sinorhizobium meliloti* GhrA (PDB ID 5UOG^*30*^) and *Pyrococcus furiosus* GhrA (PDB ID 5AOV^*33*^). The main differences lie in the conformations of the dimerization loops (residues 111-136, EcGhrA numbering) and the C-terminal section beginning at residue 296.

**Figure 1.**
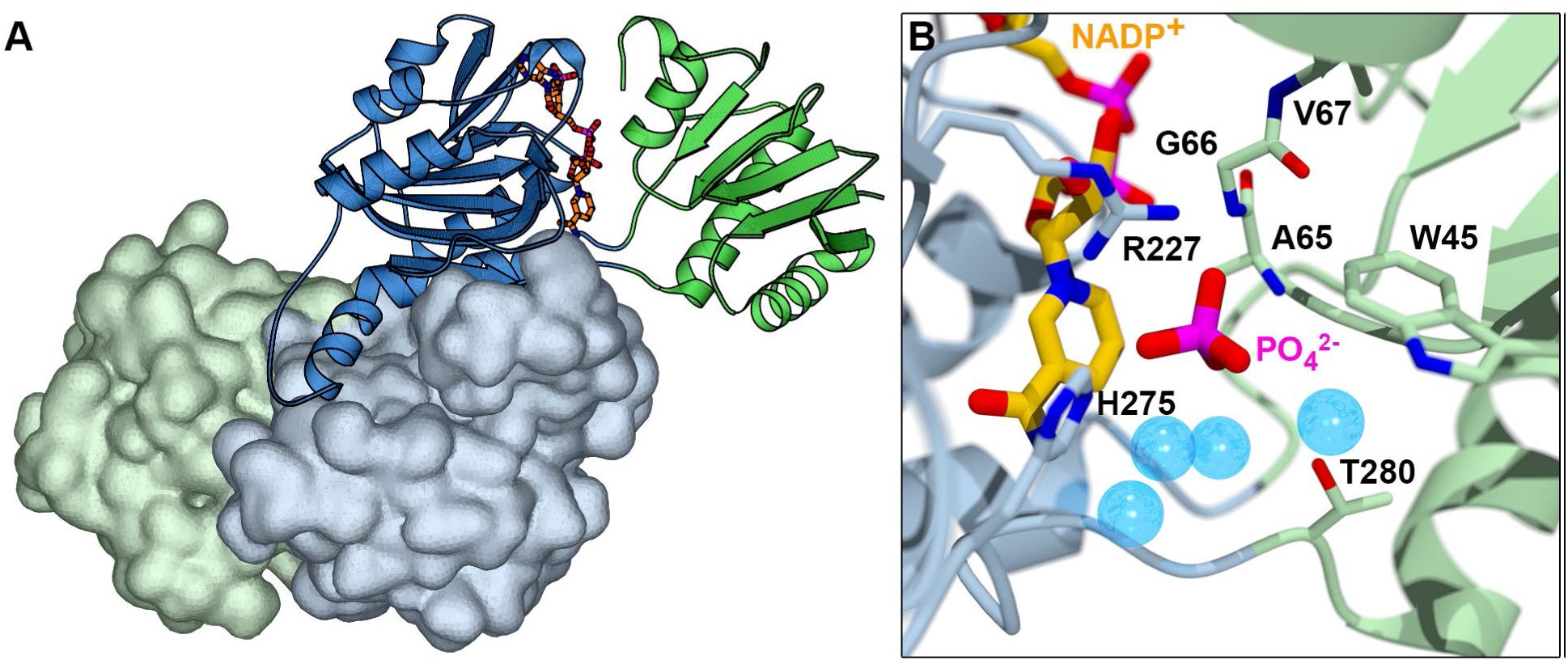
EcGhrA is active as a dimer of 35 kDa protomers as shown in (A). One protomer is shown in cartoon representation, and the other as a solvent-accessible surface. Each protomer is comprised of two domains: a nucleotide-binding domain (blue) and a substrate-binding domain (green). The active site of the EcGhrA·NADP^+^ binary complex (B) is organized exactly as observed in other GhrA homologs. The PO_4_ anion bound in the active site is from the crystallization buffer.

The question driving this work—can we generate an EcGhrA variant that does not accept α-keto acids having large side chains (*e.g*. **3** and **8**) without abrogating activity with glyoxylate (**6**)—demanded that we determine how **6, 3**, and **8** bind to the wild-type enzyme by determining structures of their ternary complexes. A challenge we encountered in this effort was that the crystallization conditions identified via high-throughput screening invariably contained high concentrations of phosphate, sulfate, or tartrate salts. We have observed that, at high concentrations, phosphate significantly inhibits EcGhrA activity (data not shown). Phosphate can be observed bound to R227 in the structure of the binary complex of EcGhrA with NADP^+^ (Figure 1B), and thus these ions compete with substrates for binding at the active site. To solve this problem, we identified a new, phosphate-free crystallization condition. The new condition contained SO_4-_ ions, but at a much lower concentration (0.2M) than the phosphate in the original condition (1.4M). When crystals grown from the phosphate-free condition were soaked with substrates as described in the Materials and Methods, the resulting crystal structures showed excellent electron density for the α-keto acid moieties of **6, 3**, and **8** (Figures 2-4).

**Figure 2.**
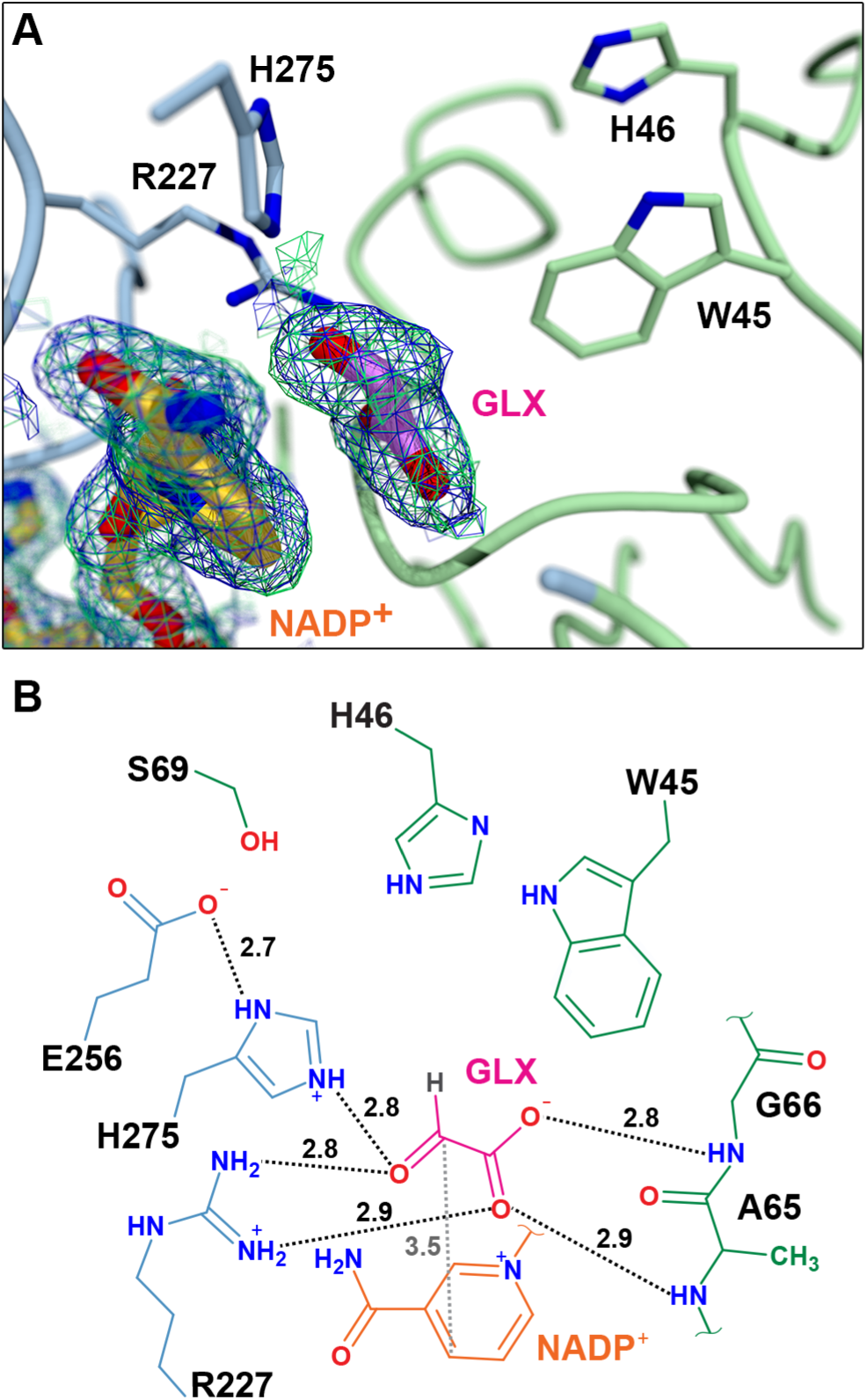
Electron density of NADP^+^ and **6** in the EcGhrA·NADP^+^·**6** ternary complex (A). The 2F_o_ -F_c_ electron density map contoured at 1.0σ is shown as a blue mesh, and the 2F_o_ -F_c_ simulated annealing composite omit map, also contoured at 1.0σ, is shown as a green mesh. The fact that the two maps are undistinguishable indicates that the electron density for the ligands is real and not the result of model bias. The schematic view of the ternary complex active site (B) shows hydrogen bonding interactions (black dotted lines) as well as the distance between the carbonyl carbon of **6** and C4 of the nicotinamide ring of NADP^+^ (grey dotted line). The distances in Ångströms are given next to each dotted line. Residues from the nucleotide-binding domain are colored blue, those from the substrate-binding domain are colored green, NADP^+^ is orange, and **6** is magenta.

The EcGhrA·NADP^+^·**6** ternary complex structure was determined to elucidate the binding mode of the preferred substrate, glyoxylate (Figure 2). Glyoxylate binds such that the carbonyl and carboxylic acid groups participate in hydrogen bonding interactions with R227. The oxygen atoms of the carboxylic acid are also able to form hydrogen bonding interactions with the mainchain amides of A65 and G66. This binding mode orients the carbonyl carbon of glyoxylate 3.5 Å from the C4 position of the nicotinamide ring of the cosubstrate. This is exactly the binding mode observed in other GhrA homologs, such as the *P. furiosus* GhrA (PDB ID 5AOV,^*33*^; Figure S2A). Unlike other GhrA homologs, EcGhrA does not appear to undergo a significant conformational change upon glyoxylate binding. NAD(H) binding does trigger a conformational change, since in the unliganded form of EcGhrA the substrate-binding domain (SBD) is completely disordered and only the nucleotide-binding domain (NBD) is visible (data not shown).

The crystal structure of the EcGhrA·NADP^+^·**3** ternary complex (Figure 3) was the most important with respect to informing our efforts to engineer EcGhrA to *not* accept **3** as a substrate. The structure shows that the keto acid moiety of **3** makes the same interactions with the enzyme as observed for glyoxylate in the EcGhrA·NADP^+^·**6** ternary complex. Surprisingly, however, the guanidinium group of **3** makes no clear contacts with the enzyme (Figure 4B). While the position of the guanidinium group of **3** is defined in the electron density, there is little or no density for the methylene carbon atoms of the side chain. This suggests that there is considerable mobility in the side chain and binding is mediated primarily by the interactions of the α-keto acid moiety with the main chain amides of A65 and G66, and the side chains of R227 and H275 (Figure 3). It should be noted that even these ternary complex structures appear to be “open” when compared to those of other EcGhrA homologs with substrates bound (Figure S2). It is conceivable that the indole nitrogen atom of W45 and/or the imidazole group of H46 could make strong contacts with the side chain of **3** if the substrate-binding domain were fully closed. The tryptophan would likely interact with the charged guanidinium group solely via cation-π interaction ^*34, 35*^, while the histidine could accept a hydrogen bond from the protonated nitrogen atoms of the guanidinium group of **3**. Based mostly on the possibility that W45 is important for binding in a presumed “closed” conformation of EcGhrA, we set out to generate the W45F and W45F/H46S variants of the enzyme.

**Figure 3.**
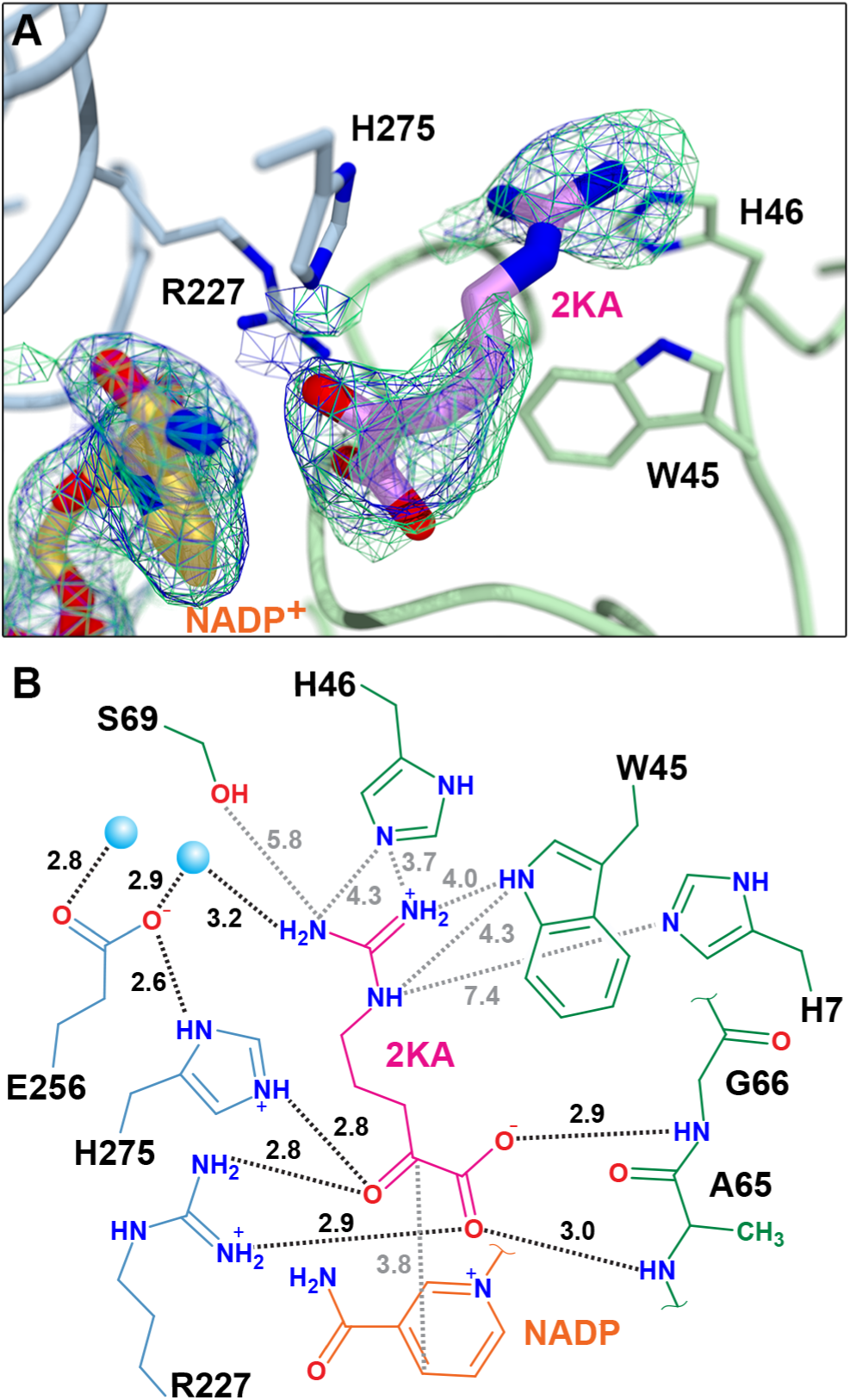
Electron density for the EcGhrA·NADP^+^·**3** ternary complex active site (A) using the same orientation as Figure 2. The 2F_o_ -F_c_ electron density map contoured at 1.0s is shown as blue mesh. The 2F_o_ -F_c_ simulated annealing composite omit map (green mesh) is also contoured at 1.0s. The schematic view (B) shows the hydrogen bonding interactions (black dotted lines) and their corresponding distances in Å. The lines do not represent non-covalent interactions, but simply indicate interesting distances. For example, the guanidinium group of **3** is relatively close (∼4.0Å) to the indole N atom of W45. With a slight movement of the side chain, **3** might participate in cation-π interactions with this residue. As in Figure 2, residues from the nucleotide- and substrate-binding domains are blue and green, respectively. The NADP^+^ is orange and the substrate **3** is magenta. The blue spheres represent water molecules.

**Figure 4.**
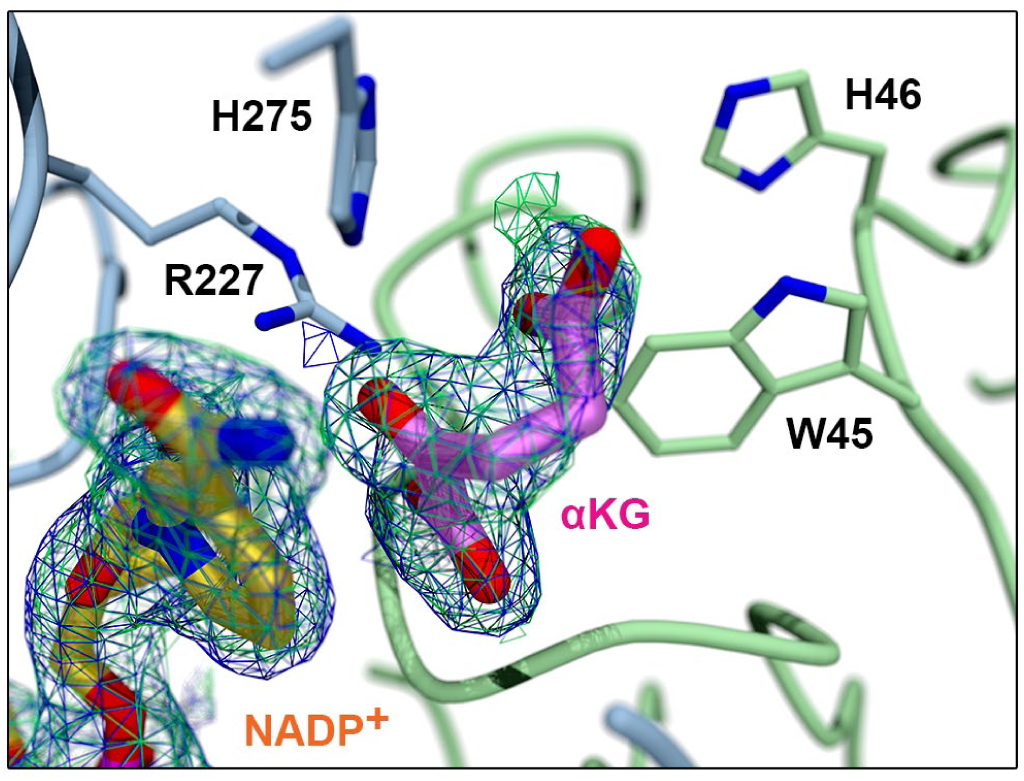
The EcGhrA·NADP^+^·**8** active site showing α-ketoglutarate bound adjacent to the cosubstrate. The 2F_o_ -F_c_ electron density map contoured at 1.0s is shown as a blue mesh, and the 2F_o_ -F_c_ simulated annealing composite omit map, also contoured at 1.0s, is shown as a green mesh. The side chain carboxylate group of **8** does not make any direct contacts with the enzyme.

The structure of the ternary complex between EcGhrA, NADP^+^, and α-ketoglutarate (**8**) shows well-defined electron density for **8** near the nicotinamide ring of NADP^+^, in the same position occupied by glyoxylate in the structure of the EcGhrA•NADP^+^•**6** complex (Figure 4). The α-keto acid moiety makes the same interactions observed in the complexes with glyoxylate and 2-ketoarginine: the carboxylic acid forms hydrogen bonding interactions with the mainchain amides of A65 and G66, the guanidinium group of R227 bridges the carboxylate and ketone groups, and the imidazole group of H275 forms a hydrogen bonding interaction with the ketone of **8**. Interestingly, there is no clear indication from the structure why the activity with **8** is so much lower than the activity against glyoxylate. Likely this difference in reactivity is due to differences in the equilibria of the keto-enol tautomerizations in these two substrates.

### Substrate specificity of EcGhrA

The kinetics of glyoxylate reduction catalyzed by wild-type EcGhrA was studied first to establish a basis of comparison with other, more thoroughly characterized, GhrA homologs. EcGhrA exhibits a strong preference for NADPH over NADH (data not shown), so all kinetic analyses were done in the presence of saturating (200 µM) NADPH. The k_cat_ value for the reaction of EcGhrA with glyoxylate (40.0 ± 23.0; Table 2, Figure S3) is similar to the values reported for the reaction of the *Homo sapiens* and *A. thaliana* homologs (27.0 and 25.5 s^-1^, respectively)^*36, 37*^. The K_M_ value for glyoxylate (2.2 ± 1.7 mM) was significantly different from those observed for the *H. sapiens* and *A. thaliana* enzymes (0.24 mM and 34 µM, respectively)^*36, 37*^. This difference may reflect differences in the structures and activities of these enzymes, or it may result from the acute substrate inhibition we observed (Figure S3; K_I_^**6**^ = 1.5 ± 1.1 mM). The simple modified Michaelis-Menten equation (Equation 1) used to fit these data resulted in poor fits, as evidenced by the large standard uncertainties on the kinetic parameters. However, more nuanced models with additional parameters either failed to converge or resulted in absurdly large standard uncertainties.

**Table 2.**
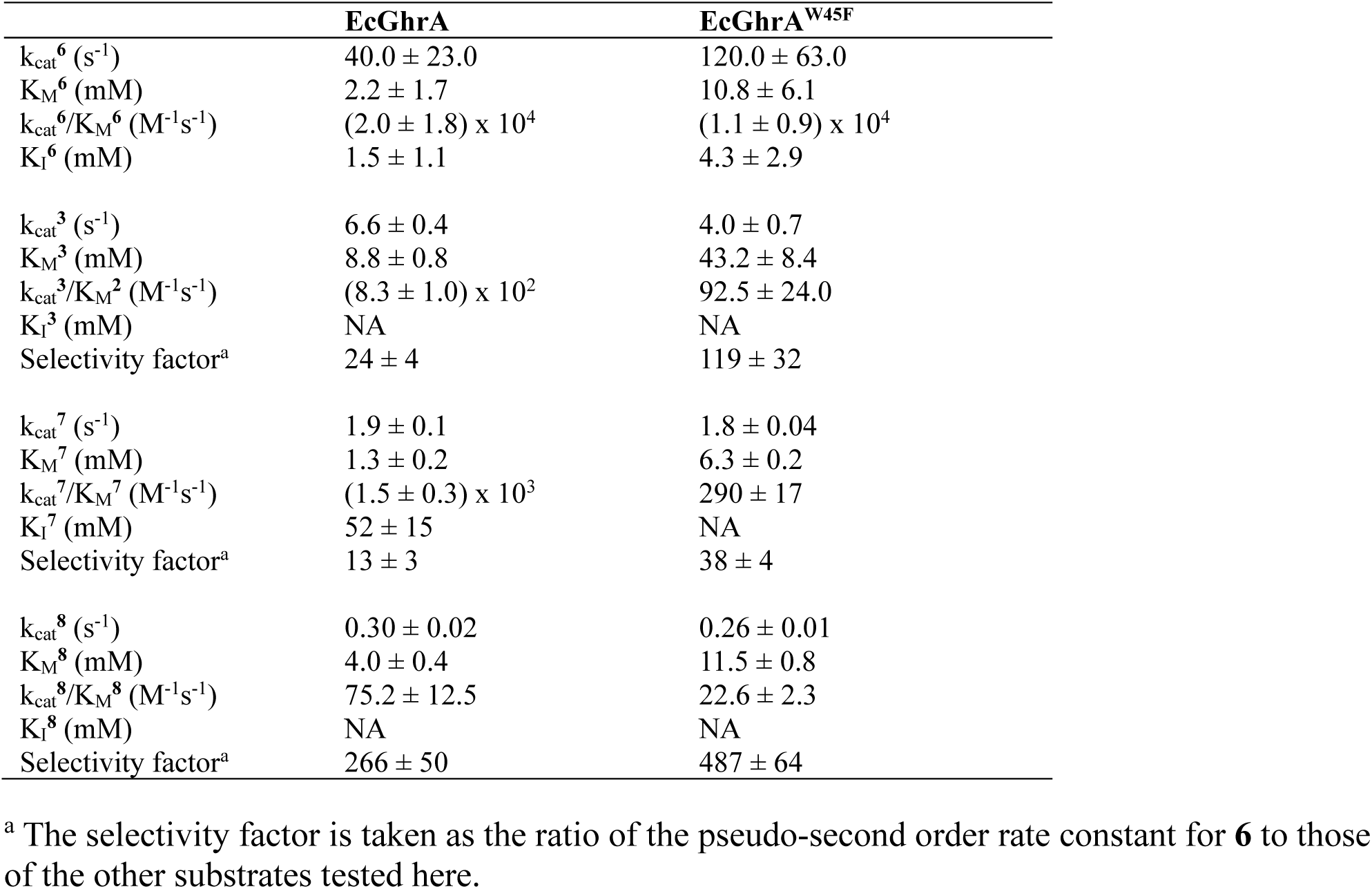
Steady state parameters of EcGhrA and EcGhrA^W45F^ with **6, 3, 7**, and **8**.

Unlike glyoxylate (**6**), 2-ketoarginine (**3**) showed no sign of substrate inhibition. The turnover number for the reaction of EcGhrA with **3** (6.6 ± 0.4 s^-1^; Table 2, Figure S4) is only 6-fold lower than that for **6** and the K_M_ value for **3** is only 4-fold higher than that for **6**. The ratio of the pseudo-second order rate constants for **6** and **3** is 24 ± 4, giving some quantitative idea of the relative preference of EcGhrA for these two substrates.

We used this ratio as a “selectivity factor” to compare the other substrates tested here (Scheme 2). Of compounds **7** – **12**, only oxaloacetate (**7**) and α-ketoglutarate (**8**) proved to be substrates for the enzyme. Oxaloacetate is nearly as efficient a substrate as **6**, and shows some evidence of substrate inhibition, though it is very weak (K_I**7**_ = 52 ± 15 mM; Table 2, Figure S5). The turnover number for **7** (1.9 ± 0.1 s^-1^) is approximately 20-fold lower than k_cat_^**6**^. The K_M_ value for **7** (1.3 ± 0.2 mM) is roughly half of **6**. The selectivity factor for oxaloacetate is only 13 ± 3, which underscores the similar catalytic efficiencies of oxaloacetate and glyoxylate. Conversely, α-ketoglutarate is a much poorer substrate, having a selectivity factor of 266 ± 50 (Table 2, Figure S6). With a K_M_ value only twice that for **6**, most of the difference between **8** and **6** comes in the turnover number. The value of k_cat_^**8**^ (0.3 ± 0.02 s^-1^) is approximately 130-fold lower than the turnover number for glyoxylate. The K_M_ values for **8** and **3** are not significantly different, which is consistent with the close agreement between the positioning of **6** and **8** in the crystal structures.

### The W45F variant of EcGhrA is more selective for glyoxylate

The W45F and H46S mutations were chosen as the most conservative changes that would still disrupt any potential interactions with substrate **3**. The W45F and W45F/H46S variants both expressed and purified similarly to the wild-type enzyme. There was no indication that mutation(s) had any effect on the stability of the enzyme. Steady state kinetic studies of both variants showed that the double mutant W45F/H46S did not behave differently from the single W45F variant. Therefore, only the W45F variant will be discussed here. EcGhrA^W45F^ exhibited a roughly 10-fold decrease in k_cat_/K_M_ for **3**, while the k_cat_/K_M_ for **6** turned out to be statistically within the range of the values for the wild-type (Table 2). Thus, the W45F mutation somewhat decreased the catalytic efficiency of the reaction with **3**, while leaving the reaction with **6** essentially untouched. The selectivity factor for **6** with respect to **3** increased 5-fold for the mutant (119 vs 24 for the wild-type). It is also important to mention that the K_M_ values for both **6** and **3** are likely overestimates, since **6** exists predominantly in a hydrated gem-diol form^*38*^, while **3** is in equilibrium with the cyclic pyrrolidine-1-amidino-2-hydroxy-2-carboxylic acid^*39*^.

The crystal structure of the single mutant showed no conformational changes of the mutated residues, or the catalytic residues R227 and H275 (Figure 5), suggesting that the substrate binding modes remain unchanged. The single W45 to Phe mutation did not disrupt the native EcGhrA reaction but did significantly reduce the reactivity with **3**. The resulting EcGhrA^W45F^ variant is thus more selective for glyoxylate and is a suitable coupling enzyme for steady state assays.

**Figure 5.**
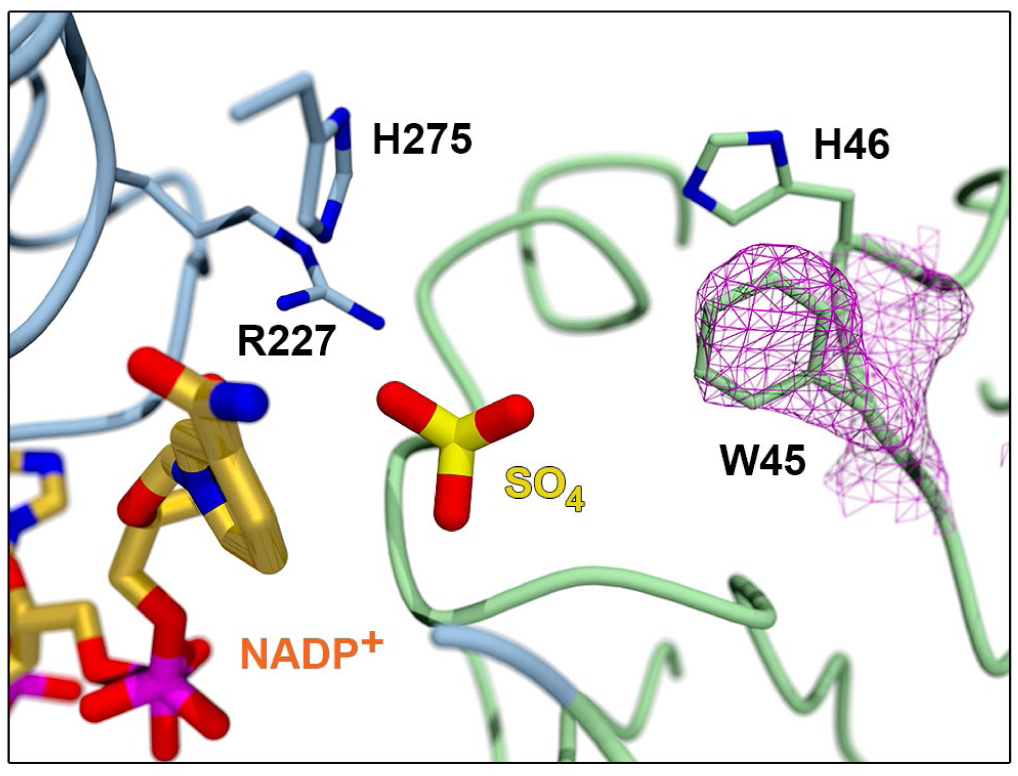
The EcGhrA^W45F^·NADP^+^ active site showing the 2F_o_ -F_c_ electron density map around the W45F mutation (magenta mesh, contoured at 1.0s). In the absence of substrate, the sulfate anion from the crystallization buffer binds near the cosubstrate.

## ABBREVIATIONS

GLX: glyoxylate
2KA: 2-ketoarginine
EcGhrA: *E. coli* glyoxylate reductase
L-End: L-enduracididine
OA: oxaloacetate
αKG: α-ketoglutarate

## TABLE OF CONTENTS USE ONLY

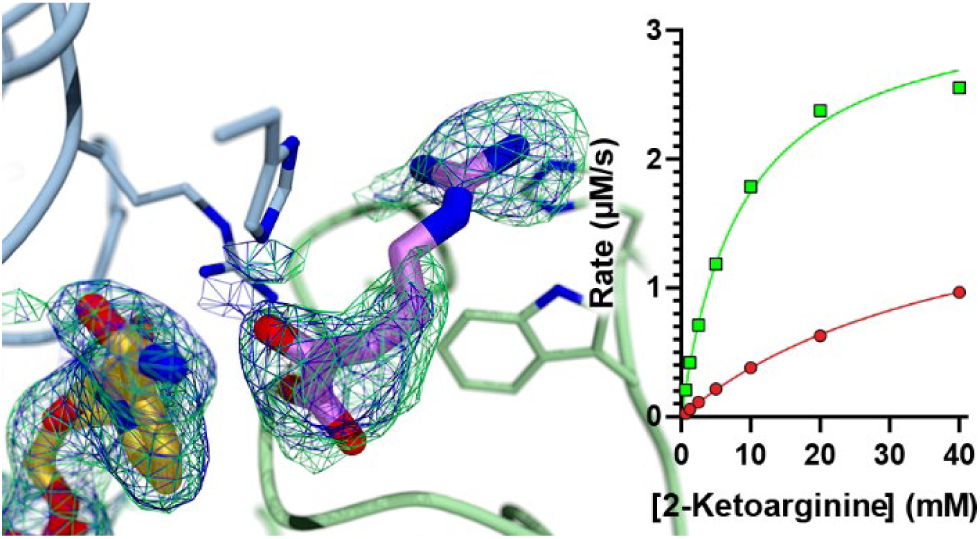

