## Supplemental Information for "Engineering a more specific *E. coli* glyoxylate/hydroxypyruvate reductase for coupled steady state kinetics assays"

### Figure S1. NMR spectra of enzymatically produced 3

(A) ^1^H NMR spectrum of **3** in D_2_O (B) ^13^C NMR spectrum of **3** in D_2_O; (C) HSQC-multiplicity edited spectrum of **3** in D_2_O

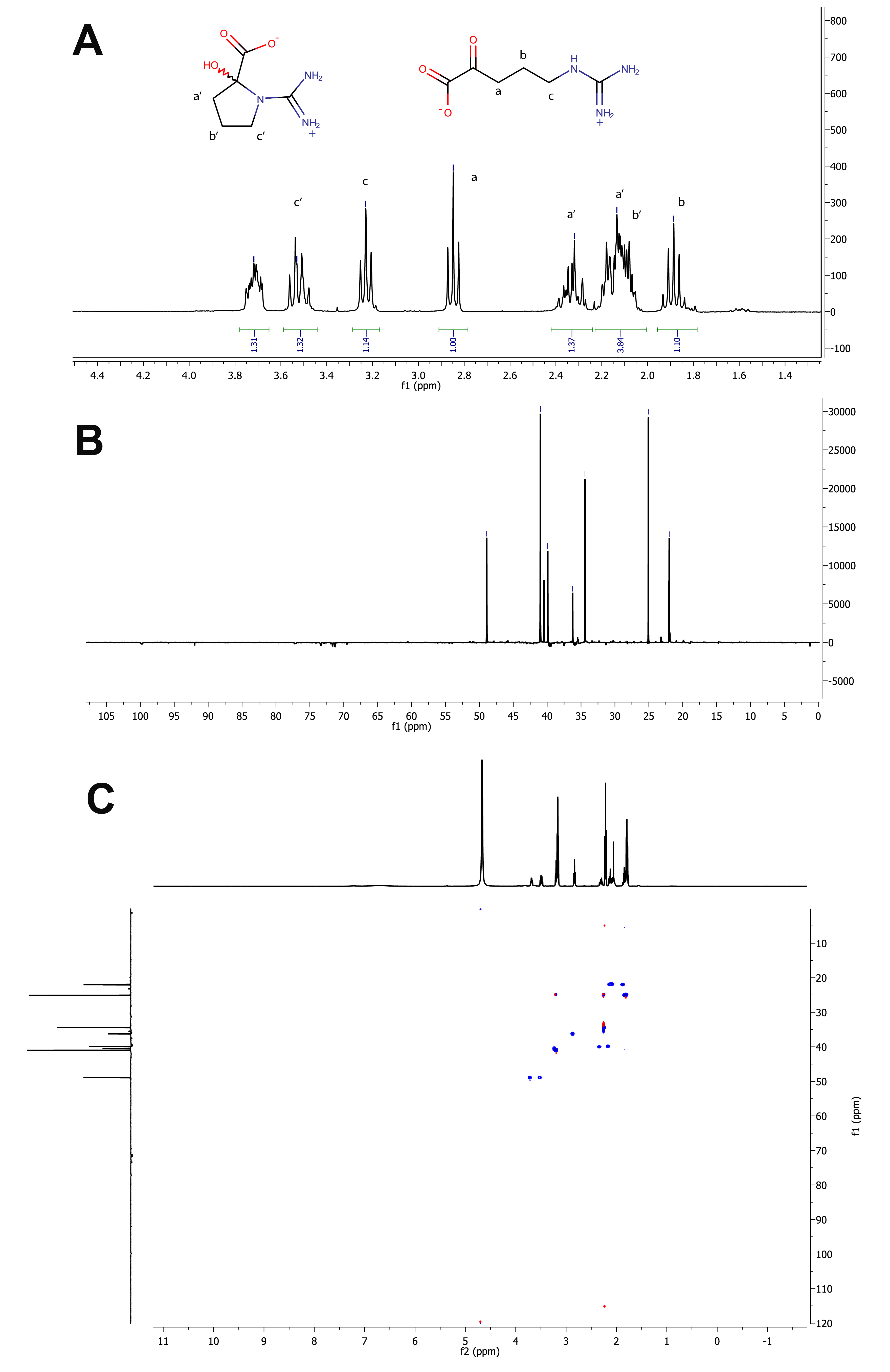

### Figure S2. Michaelis-Menten plot of wild-type EcGhrA (A) and EcGhrA^W45F^ (B) with 6

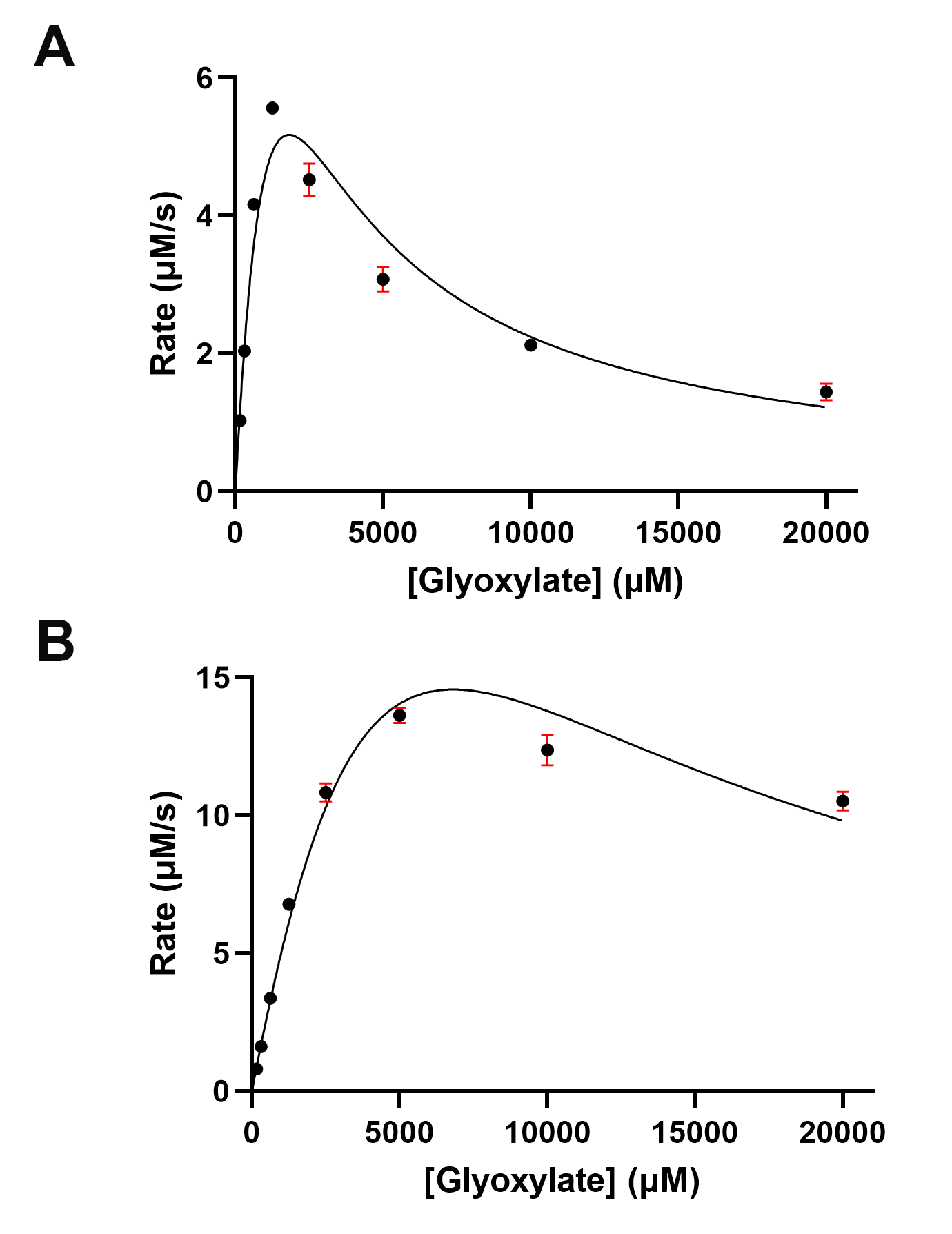

### Figure S3. Michaelis-Menten plot of wild-type EcGhrA (A) and EcGhrA^W45F^ (B) with 3

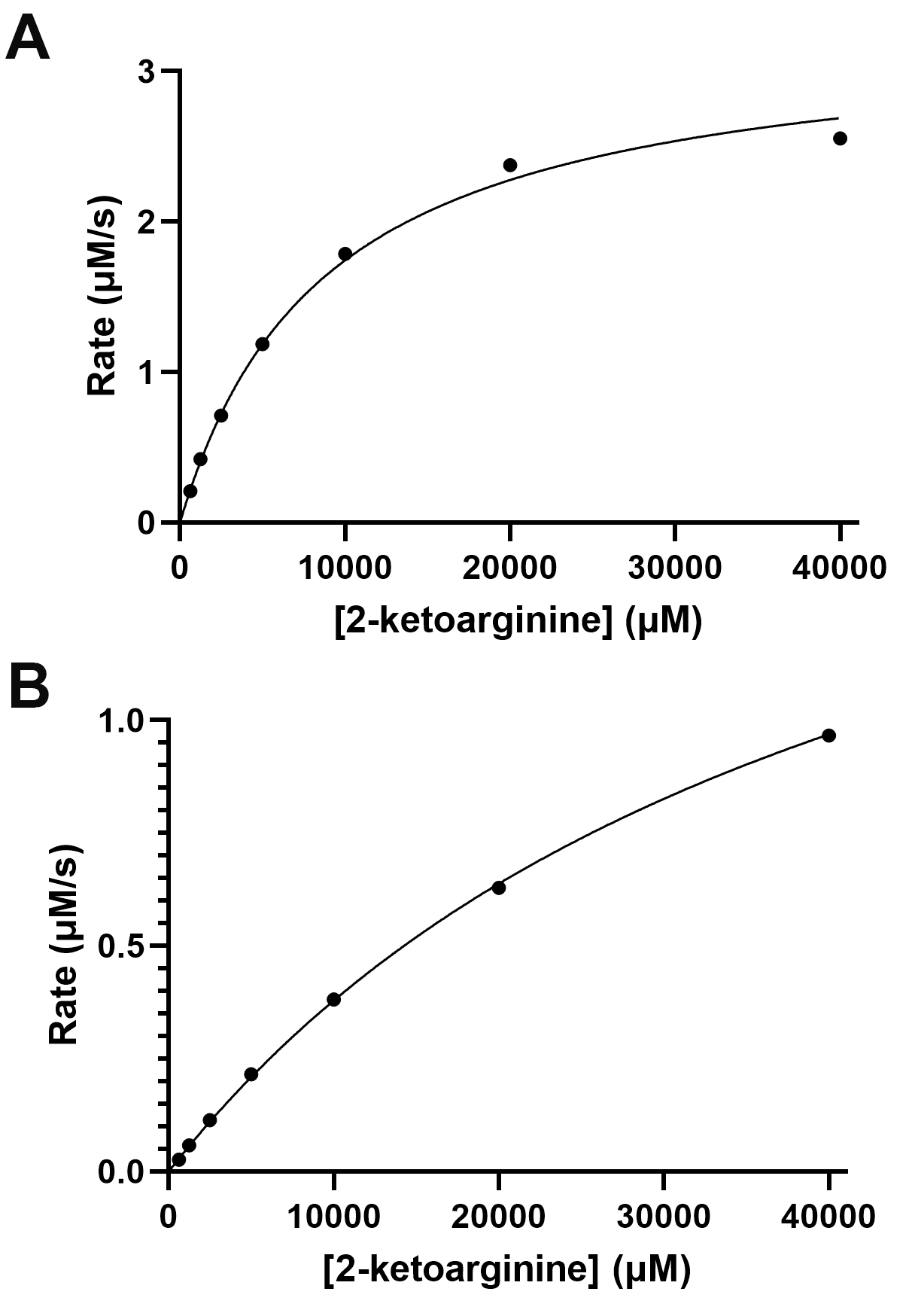

### Figure S4. Michaelis-Menten plot of wild-type EcGhrA (A) and EcGhrA^W45F^ (B) with 7

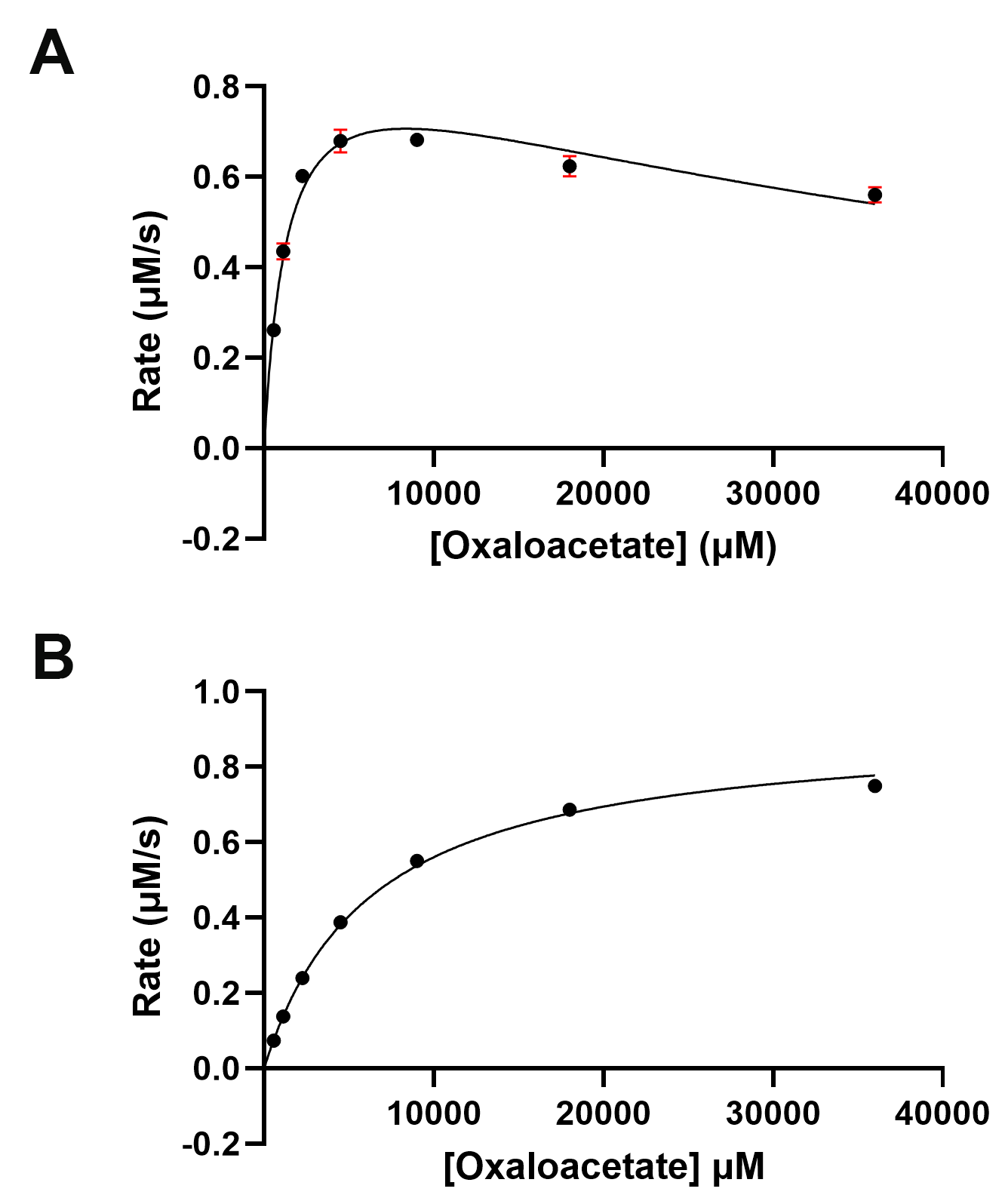

### Figure S5. Michaelis-Menten plot of wild-type EcGhrA (A) and EcGhrA^W45F^ (B) with 8

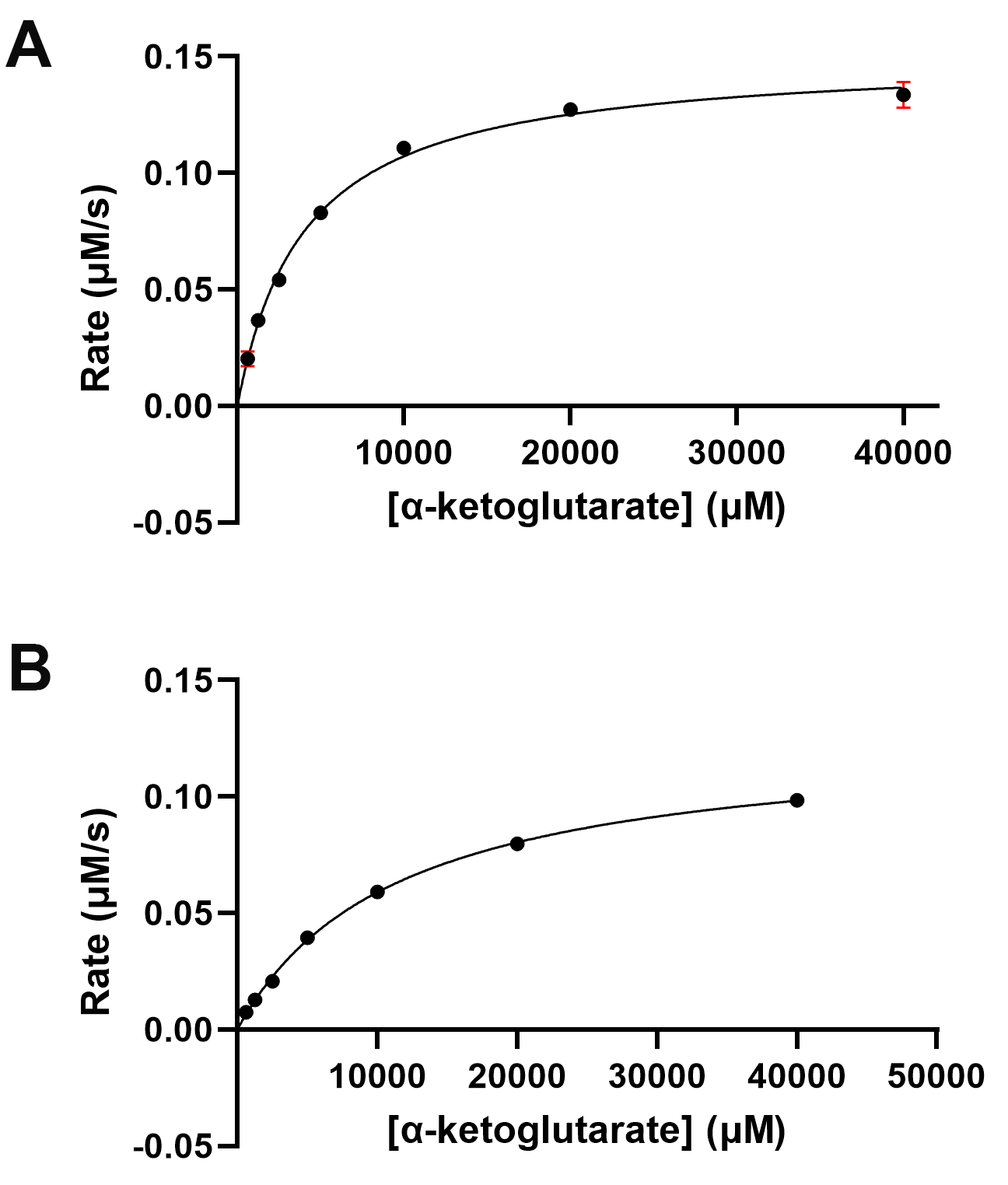

### Figure S6. Overlay of EcGhrA complexed with 3 (green) and P. furiosus glyoxylate/hydroxypyruvate reductase complexed with 6 (PDB entry: 5AOV) (magenta).

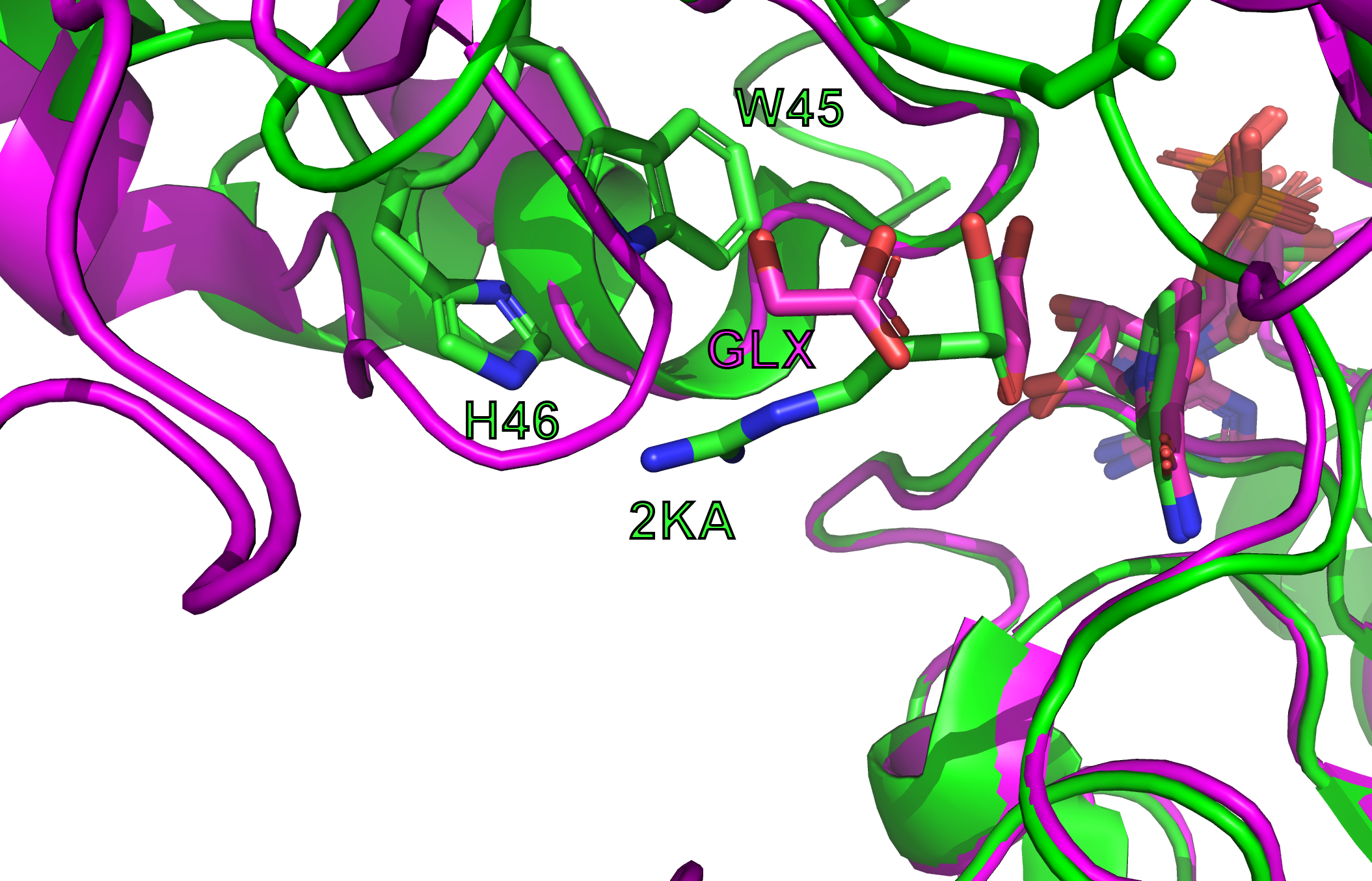
